# Flagellin-mediated TLR5 activation enhances innate immune responses in healthy and diseased human airway epithelium

**DOI:** 10.1101/2025.11.06.686934

**Authors:** Xing Li, Delphine Cayet, Yasmine Zeroual, Itziar Sanjuán-García, Amélie Bonnefond, Mehdi Derhourhi, Mireille Caul-Futy, Christine Caul-Futy, Tom van der Poll, Christophe Carnoy, Mara Baldry, Samuel Constant, Jean-Claude Sirard

## Abstract

Bacterial pneumonia poses a significant challenge to public health, often leading to antibiotic treatment failure. Enhancing innate immunity represents a promising adjunctive strategy to conventional antibiotic therapy. Bacterial flagellin, a Toll-like receptor 5 (TLR5) agonist, has been shown to stimulate innate immune defenses when delivered via the respiratory route, demonstrating efficacy in both preventing and treating bacterial pneumonia in murine models. This protective effect is primarily mediated through TLR5-driven activation of airway epithelial cells. This study aimed to characterize the immunomodulatory effects of flagellin on human primary respiratory epithelium. Using the MucilAir™ air-liquid interface model and RNA sequencing, we demonstrated that apical administration of flagellin induced robust immune responses in airway epithelium derived from healthy individuals, as well as patients with chronic obstructive pulmonary disease (COPD) and cystic fibrosis (CF). TLR5-mediated epithelial signaling triggered key immune-related pathways, including cytokine production, leukocyte chemotaxis, neutrophil recruitment, and antimicrobial defense, with strong commonalities across healthy and diseased airway epithelia. Furthermore, we demonstrated that flagellin effectively activated epithelial immune responses even in the presence of the bacteria *Pseudomonas aeruginosa* or *Streptococcus pneumoniae*. However, epithelial activation alone was insufficient to directly limit bacterial colonization or replication, highlighting the potential role of epithelial-immune cell interactions in achieving effective bacterial clearance. These findings support TLR5 activation as a promising therapeutic strategy to enhance host defense mechanisms and improve treatment outcomes for bacterial pneumonia in both healthy individuals and patients with COPD or CF.

## Introduction

Innate immunity serves as the first line of defense against infectious agents, providing a ubiquitous and broad-spectrum response to pathogenic microorganisms (1). The activation of innate immunity relies on the recognition of danger signals or microbe-associated molecular patterns (MAMPs), which originate from microorganisms and are detected by pattern recognition receptors (PRRs). Among these, Toll-like receptors (TLRs) represent the most extensively studied PRR family and are highly expressed by sentinel cells, including immune and structural cells (2). Given its critical role in host defense, the targeted activation of innate immunity is of significant interest for the prevention and treatment of infections.

Surface exposed TLR5 senses bacterial flagellin, the major structural protein of flagella (3–5). The flagellin FliC from *Salmonella enterica* serovar Typhimurium, that represents the archetype TLR5 agonist, is composed of four distinct domains: D0 and D1 which form the inner core of the flagellar filament and contain the molecular motifs essential for TLR5 recognition, and D2 and D3, which are highly variable and exposed on the filament surface (4). These structural features play a critical role in modulating immune responses upon host-pathogen interactions. Previous studies have demonstrated that recombinant flagellin variants, such as FliC_Δ174–400_ (also known as FLAMOD), which lack the hypervariable D2 and D3 domains, retain their TLR5-activating properties and effectively stimulate mucosal immunity (6–8). At mucosal surfaces, epithelial cells serve as the primary sentinels that detect flagellin and orchestrate immune responses. Intestinal, respiratory, and urogenital epithelial cells exhibit robust responsiveness to flagellin, leading to activation of downstream pro-inflammatory pathways (reviewed in (4)). Following respiratory administration of flagellin, TLR5 engagement on epithelial cells triggers the production of key mediators, including chemokines such as CXCL8 (IL-8) and CCL20, which facilitate the recruitment of myeloid and lymphoid immune cells to the airways (4). In addition to chemokine production, flagellin stimulates epithelial cells to secrete cytokines and growth factors such as IL-6, IL-1β, and granulocyte colony-stimulating factor (G-CSF), which further promote the activation, proliferation, and survival of recruited immune cells. Beyond their role in immune signaling, epithelial cells also possess direct antimicrobial capabilities. Upon flagellin stimulation, they upregulate the expression of antimicrobial peptides (AMPs) and enhance barrier integrity, providing a crucial first line of defense against invading pathogens. Several reports have demonstrated the potent anti-infectious properties of flagellin in murine models of respiratory infection. The administration of flagellin via the respiratory route, either intranasally or intratracheally, has been shown to confer protection against respiratory pathogens such as *Streptococcus pneumoniae* and *Pseudomonas aeruginosa* (9–11). The protective effect against *S. pneumoniae* is observed not only when flagellin is used prophylactically but also when administered therapeutically in conjunction with antibiotics, enhancing bacterial clearance and reducing disease severity (7, 8, 12–14). The ability of flagellin to modulate both immune and defense mechanisms highlights its potential as an immunostimulatory agent for respiratory interventions.

Human primary airway epithelial cells and respiratory cell lines have been shown to mount a robust response to TLR5 stimulation, leading to the production of a diverse array of immune mediators (15–26). Respiratory diseases, including chronic obstructive pulmonary disease (COPD) and cystic fibrosis (CF), are characterized by impaired epithelial innate immunity and defense mechanisms, thus increasing susceptibility to respiratory infections (26–28). These conditions involve structural and functional epithelial alterations that weaken protective immune responses. In COPD, airway epithelial cells exhibit significantly downregulated TLR5 expression, reducing responsiveness to flagellin stimulation (18). In contrast, in CF airway epithelium, flagellin stimulation triggers exacerbated pro-inflammatory responses (16, 29). This crosstalk between epithelial innate immunity and chronic respiratory disease pathophysiology highlights the need for further investigation and benchmarking of flagellin-based interventions.

In this study, we compared flagellin-mediated responses in a human model using primary respiratory epithelium derived from healthy individuals and patients with COPD or CF. To this end, we characterized the transcriptional response via RNA sequencing (RNA-seq) to elucidate the molecular mechanisms underlying epithelial activation. This approach allowed us to identify key signaling pathways and innate immune defense mechanisms that exhibit strong commonalities across healthy, CF, and COPD airway epithelium. Finally, we assessed the capacity of flagellin to activate epithelial cells from healthy individuals in the context of bacterial infection with *Pseudomonas aeruginosa* or *Streptococcus pneumoniae*.

## Results

### Flagellin stimulates innate immune defense signatures in respiratory epithelium of healthy subjects

As TLR5 signaling is strongly associated with transcriptional activity, we performed RNA-seq analysis to investigate the innate response to flagellin in human airway epithelium. The recombinant flagellin FliC_Δ174-400_ 0.7 µg in 10 μl of PBS (*i.e.,* 2.1 µg/cm^2^) was deposited at the air interface of MucilAir^TM^ epithelium from a pool of 14 healthy subjects. As a mock treatment, PBS was applied on the epithelium. After a 4 h stimulation, total RNA was extracted and processed for RNA-seq analysis (**Figure 1**). More than 3,200 DEG (FDR-adjusted p < 0.05) were observed in response to flagellin compared to PBS-treated epithelium (**Supplementary Figure 1 and Supplementary File 1**). When focusing on genes with log_2_|fold-change| (log_2_|FC|)≥1, the analysis identified 843 up- and 283 down-regulated genes. The most substantially upregulated genes included *CCL4, CCL20, CSF3, CXCL2, CXCL5, CXCL6, CXCL8, DEFB4A, IDO1, IL1B, IL17C, IL23A, MUC2, MUC5AC, RELN, TFF1, TNFAIP6*, while the genes with the highest decrease in transcript levels were *APELA, CLDN8, PTGFR, TACR1*, and *TNFSF4* (**Figure 1A**). To better characterize the relation between DEG and biological response, functional annotation was performed using GSEA (**Figure 1B and Supplementary File 2**). Enriched pathways were found to be related to immune cell activity, activation and migration pathways (cytokine activation, leucocyte chemotaxis, neutrophil migration) and antimicrobial/antibacterial defenses. To validate the transcriptomic analysis, a new set of experiments was conducted and analyzed by RT-qPCR on selected up- and down-regulated genes (**Figure 1C**). The mRNA levels of the *CSF3, DEFB4A, IL17C* and *TNFAIP6* genes encoding the neutrophil stimulating factor G-CSF, the antimicrobial peptide beta-defensin 2, the epithelial cytokine IL-17C and the extracellular regulator of inflammation TNFAIP6 were increased 18- to 420-fold at 4h by flagellin stimulation. In contrast, mRNA levels of *CLDN24* and *TACR1* genes encoding the tight junction component Claudin 24 and the neurokinin 1 receptor were globally decreased 3- to 12-fold by flagellin treatment. Kinetic analysis revealed a peak in stimulatory activity at 2 h following flagellin exposure, with substantial gene expression modulation still evident at 4 h (**Supplementary Figure 2A**). Finally, dose-response analysis estimated the ED50 (the dose required to elicit half-maximal biological activity) of flagellin on airway epithelium to be 0.006 µg/cm² (equivalent to 0.2 picomole/cm²) (**Supplementary Figure 2B**). In conclusion, human respiratory epithelial cells from healthy individuals readily respond to the application of flagellin at the air interface.

**Figure 1.**
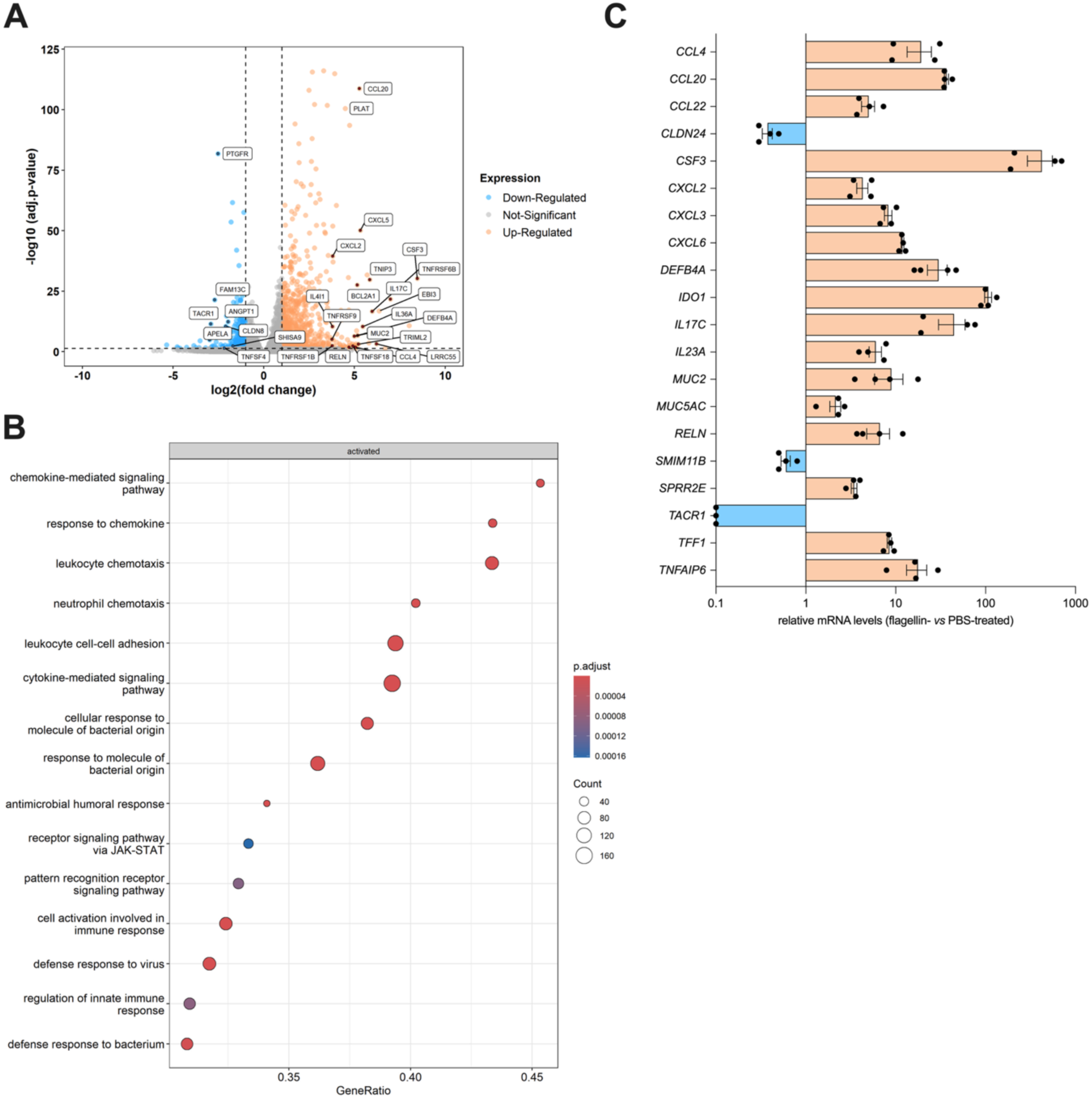
Transcriptional response to flagellin of respiratory epithelium from healthy donors. Flagellin FliC_Δ174-400_ (0.7 µg in 10 µL) or the diluent solution (PBS) was applied at the air interface of MucilAir™ cells. After 4 h incubation, RNA was processed for RNA-seq analysis (n=3 per condition) or gene expression analysis by RT-qPCR (n=4 per condition). **(A)** Volcano plot of differentially-expressed genes in epithelium between flagellin- and PBS-treated conditions. **(B)** GSEA analysis. Enriched pathways related to innate and adaptive immunity in response to flagellin were listed according to gene ratio, gene numbers and *p*-adjusted values. **(C)** Validation of transcriptional response by RT-qPCR. mRNA levels were normalized for each gene house-keeping genes and the condition treated with PBS was set arbitrarily at the value of 1. Results are representative of 2 experiments. Mann-Whitney statistical analysis demonstrates *p<0.05* for each gene.

### Transcriptomic analysis of flagellin response in airway epithelium from COPD and CF patients reveals significant similarities to that of healthy epithelium

To define whether TLR5 signaling is altered in pathological conditions, flagellin was added at 2.1 µg/cm^2^ on epithelial cells from patients with COPD or CF. After 4 h incubation, RNA-seq was performed and normalized to the PBS treatment (**Figure 2 and Supplementary File 1**). Similarly to healthy subject epithelium, analysis revealed more than 5,000 and 2,500 DEG in COPD and CF epithelium, respectively (**Supplementary Figure 1**). Analysis of genes with log_2_|FC|≥1 compared to cells from healthy subjects revealed 851 and 930 upregulated genes, and 465 and 401 downregulated genes, in COPD and CF conditions, respectively. In **Figure 2A-B**, the volcano plots highlighted the same genes as for the healthy condition suggesting a similar pattern of regulation. The GSEA analysis further showed that the enriched pathways in COPD and CF epithelium are also related to innate signaling, immune cell chemotaxis and activation, and antimicrobial defense response (**Figure 2C and Supplementary File 2**). Experiments with RT-qPCR validated the RNA-seq observations (**Figure 2D-E**). We next conducted a comparative analysis of flagellin-stimulated airway epithelium derived from healthy individuals, as well as patients with COPD or CF (**Supplementary Figure 3**). The Venn diagram identified 539 genes common to all conditions, representing 40-45% of DEG with log_2_|FC|≥1. Circos plots further illustrated that most genes and biological pathways are shared across the three conditions. Notably, the predominant pathways in response to flagellin stimulation were associated with the innate immune response, cytokine signaling in the immune system, and regulation of immune effector processes. Collectively, these findings highlight the conserved and robust immune-activating properties of flagellin in human airway epithelial cells, irrespective of health or disease status.

**Figure 2.**
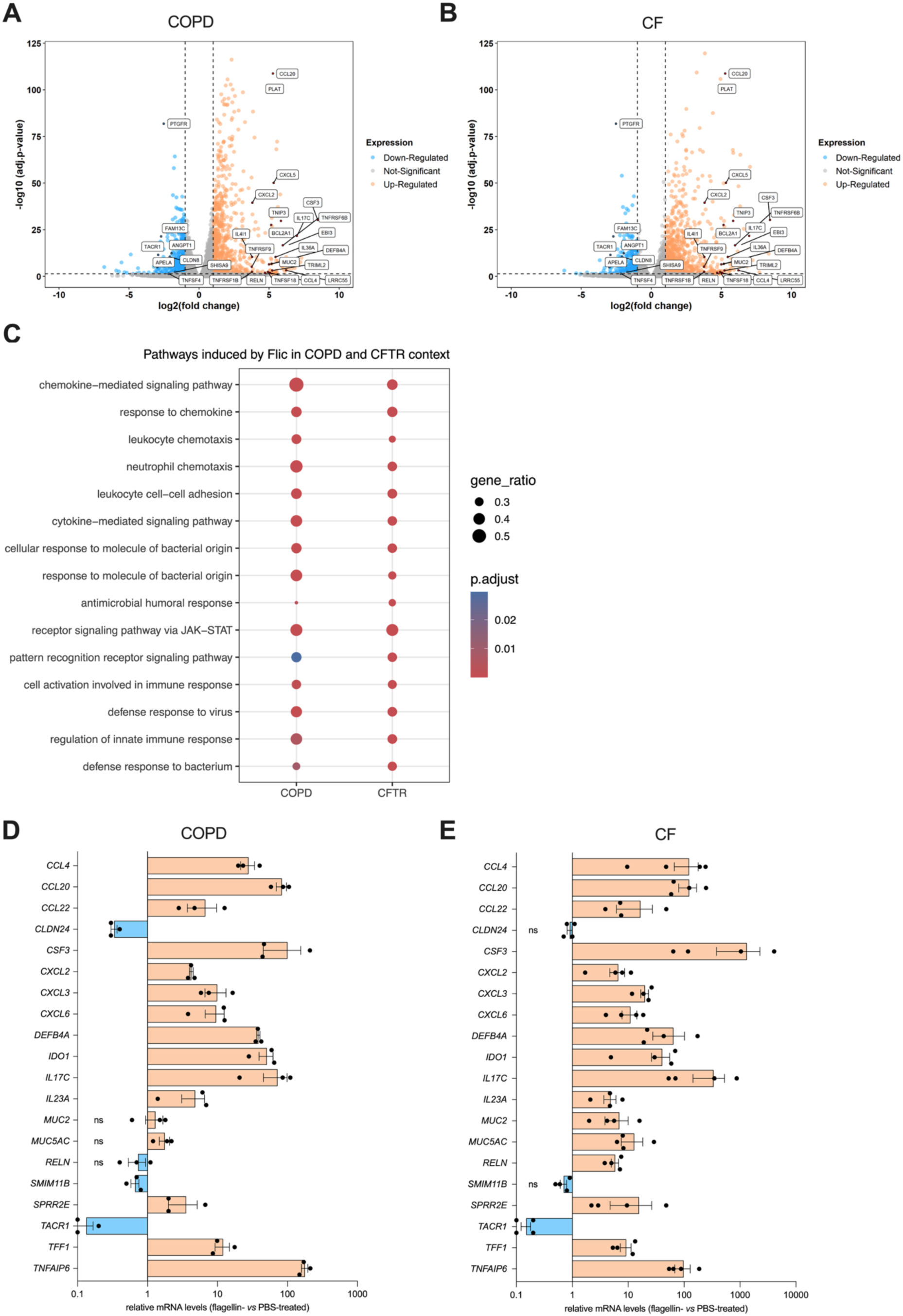
Transcriptional response to flagellin of respiratory epithelium from COPD and CF <u>patients</u>. Flagellin FliC_Δ174-400_ (0.7 µg in 10 µL) or the diluent solution (PBS) was applied at the air interface of MucilAir™ cells from COPD patients **(A,C,D)** or CF patients harboring the *CFTR*<u>Δ*508*</u> mutation **(B,C,E)**. After 4 h incubation, RNA was processed for RNA-seq analysis (n=3-4 per condition) or gene expression analysis by RT-qPCR (n=3-4 per condition). **(A-B)** Volcano plot of differentially-expressed genes in epithelium between flagellin- and PBS-treated conditions. **(C)** GSEA analysis. Enriched pathways in response to flagellin were listed according to gene ratio, gene numbers, and P-adjusted values. **(D-E)** Validation of transcriptional response by RT-qPCR. mRNA levels were normalized for each gene house-keeping genes and the condition treated with PBS was set arbitrarily at the value of 1. Mann-Whitney statistical analysis demonstrates *p<0.05* for most genes; ns stands for non-significant.

### Flagellin still induces a robust response in healthy human respiratory epithelium during bacterial infection

To evaluate the potential of flagellin as a therapeutic strategy for infections, we next investigated whether the immune-stimulating effects remained active when airway epithelium was exposed to pneumonia-associated bacterial pathogens. To this end, airway epithelium derived from healthy individuals were infected at the apical compartment with a precisely calibrated inoculum of 50 CFU of the respiratory pathogen *P. aeruginosa*. In two independent experiments, cells were incubated for 17 h following inoculation in the first experiment and 16 h in the second. Subsequently, the apical compartment was treated with either flagellin or PBS (control). Two hours post-treatment, airway epithelia were processed for downstream analyses (**Figure 3**). Bacterial quantification post-infection and treatment revealed a range of 70 to 1,770 CFU per filter in the first experiment and approximately 4×10⁴ to 10⁶ CFU in the second, highlighting variability across biological replicates (**Figure 3A**). Despite variability, these findings confirmed successful colonization of the apical compartment by *P. aeruginosa* and indicate that flagellin treatment did not significantly alter bacterial proliferation. Transcriptional analysis of selected genes demonstrated significant upregulation in flagellin-treated epithelium compared to PBS-treated controls (**Figure 3B-I**). Specifically, mRNA levels of immune-related genes *CCL20*, *CSF3*, *CXCL2*, *CXCL8*, *DEFB4A*, *IL1B*, *IL17C*, and *TNFAIP6* were significantly elevated following flagellin exposure in infected epithelial cells. These findings suggest that *P. aeruginosa* infection does not impair flagellin-induced pro-inflammatory responses in respiratory epithelial cells derived from healthy individuals.

**Figure 3.**
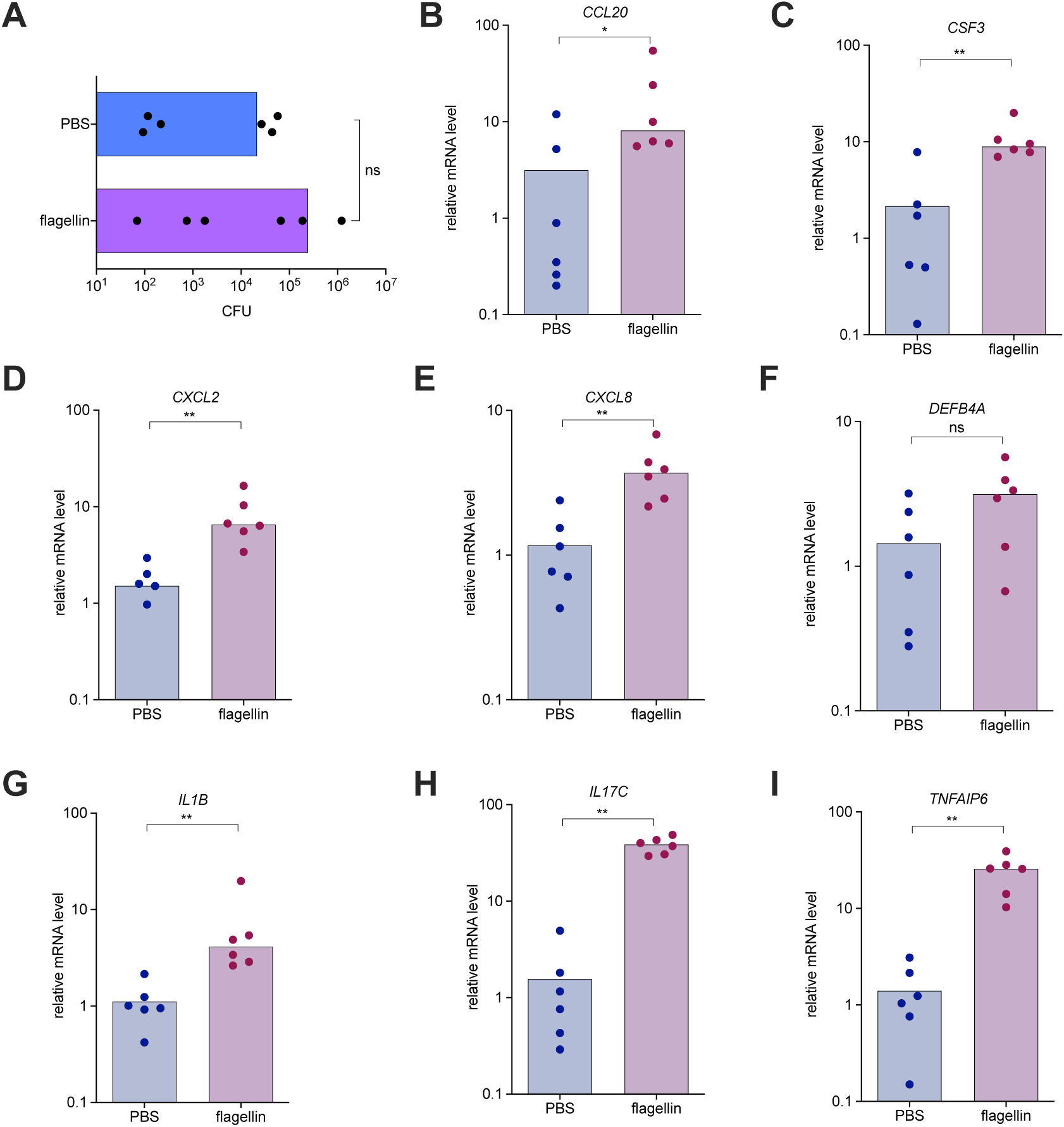
Response of airway epithelium to flagellin during *P. aeruginosa* infection. MucilAir™ respiratory epithelial cells from healthy donors (n=6 per group) were infected with *P. aeruginosa* by applying 20 µL of a calibrated bacterial suspension (16 or 52 CFU) to the apical compartment. Following 16–17 h incubation at 37°C with CO₂, the infected cells were stimulated with flagellin FliC_Δ174-400_ (0.7 µg in 10 µL) or PBS (control) applied to the apical air interface. The apical compartment was then washed with 200 µL PBS to assess bacterial growth **(A)**, and total RNA was extracted for RT-qPCR analysis **(B-I)**. **(A)** Bacterial Colonization. The apical washes were serially diluted and plated on agar to determine CFU numbers. **(B–I)** Relative transcript levels. mRNA levels were normalized to housekeeping genes and to the PBS-treated infected condition, which was arbitrarily set to 1. Statistical analysis was performed using Mann-Whitney test.

Similar experiments were conducted with epithelial cells that were infected at the apical compartment with 3×10^4^ *S. pneumoniae*. In this setting, the apical compartment was treated 1 h post-infection with either flagellin or vehicle (control). Moreover, amoxicillin was added in the basal compartment to mimic the standard of care. Twenty-four hours post-infection, cells were processed for downstream analyses (**Figure 4**). Bacterial quantification revealed a median of 2.27×10^5^ CFU in vehicle-treated controls (**Figure 4A**). Notably, the flagellin treatment did not significantly alter bacterial load (median of 3.09×10^5^ CFU), whereas amoxicillin alone and the combination of flagellin with amoxicillin effectively reduced bacterial growth by 382_to_3635-fold. Further investigation of host immune responses revealed that flagellin treatment significantly modulated β-defensin 2 and IL-8 production in *S. pneumoniae*-infected epithelial cells compared to vehicle-treated uninfected control epithelium (**Figure 4B-C**). These data indicate that *S. pneumoniae* infection does not compromise the ability of respiratory epithelial cells to mount flagellin-induced immune responses. Collectively, our findings provide strong evidence that respiratory epithelium infected with either Gram-negative or Gram-positive bacteria retains a robust capacity to respond to TLR5 activation, highlighting the potential of flagellin to be applied as an immunomodulatory agent for next-generation pneumonia treatments.

**Figure 4.**
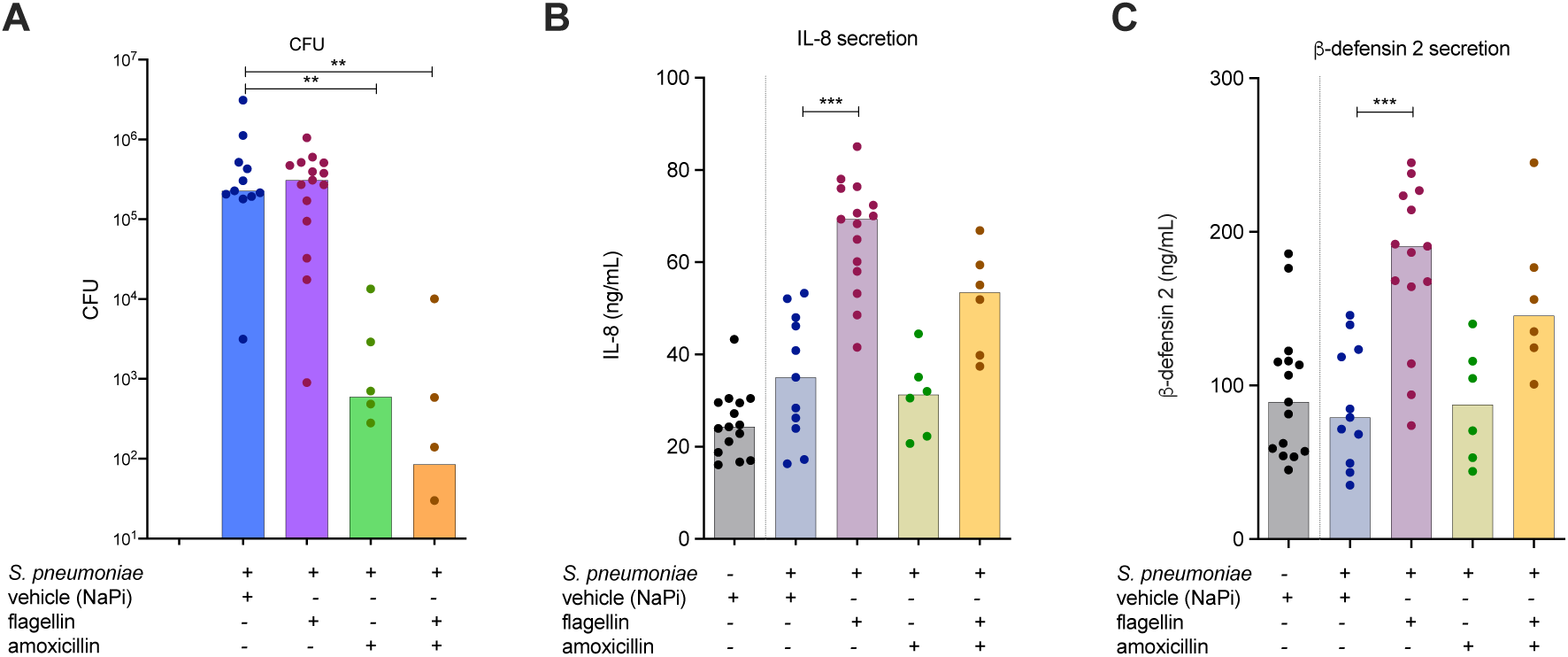
Response of airway epithelium to flagellin in the context of *S. pneumoniae* infection. MucilAir™ respiratory epithelial cells from healthy donors (n=6-15 per group) were infected with *S. pneumoniae* by applying 20 µL of a calibrated bacterial suspension (3×10^4^ CFU) to the apical compartment. Following 1 h incubation at 37°C with CO₂, the infected cells were stimulated for 23 h with flagellin FLAMOD (0.1 µg in 10 µL) or vehicle NaPi buffer (control) applied to the apical air interface or amoxicillin added in the basal medium at 1.5 µg/ml. Then, the apical compartment was washed with 200 µL of PBS to assess bacterial growth **(A)**. Bacterial colonization was measured on apical washes that were serially diluted and plated on agar to determine CFU numbers. Basal medium was used to measure IL-8 **(B)** and beta-defensin 2 **(C)** secretion by ELISA. Statistical analysis was performed using one-way ANOVA Kruskal-Wallis test compared to the infected epithelium treated with vehicle.

## Discussion

This study demonstrates the potent immunomodulatory activity of flagellin in primary human respiratory epithelium. Notably, flagellin elicited a consistent immune response not only in epithelial cells from healthy individuals but also in those derived from patients with chronic respiratory diseases such as COPD and CF. Transcriptomic analysis via RNA-seq revealed a highly conserved activation pattern across all three types of donors, highlighting the robustness and reproducibility of TLR5-induced immune stimulation. Furthermore, our findings show that flagellin-mediated pro-inflammatory signaling remains effective even in the presence of *P. aeruginosa* or *S. pneumoniae* infection, suggesting that bacterial colonization does not impair TLR5-driven immune responses. These results provide compelling evidence that picomolar dose of flagellin can rapidly activate epithelial defense mechanisms irrespective of underlying respiratory pathology. In conclusion, this study establishes a strong rationale for the development of flagellin-based biotherapeutics, particularly as an inhaled immunomodulator for the treatment and prevention of bacterial pneumonia.

Differences in TLR5 expression and activation in the airway epithelium have been reported for both COPD and CF contexts, leading to distinct responses to flagellin stimulation (16, 18, 29). In COPD associated with smoking, TLR5 expression is downregulated in the airway epithelium, rendering it unresponsive to flagellin stimulation compared to healthy airways, and leading to reduced production of key downstream immune effectors, including IL-6 and IL-8 (18). In contrast, CF airway epithelium has been shown to exhibit an exacerbated pro-inflammatory response upon flagellin-mediated activation (16, 29). However, our study did not observe heightened and decreased inflammatory responses in CF and COPD airway epithelium, respectively, relative to healthy controls. The use of recombinant flagellin from *S.* Typhimurium with deleted immunogenic domains or the effective dose compared to the referenced CF studies used *P. aeruginosa* flagellin could account for the discrepancy in findings. Interestingly, we found comparable baseline levels of *TLR5* mRNA (transcripts per million) and genes coding the TLR signaling machinery (*MYD88, IRAK2/3, TNFAIP3, IRF3/5/7, TRAF2/3/6, NFKBIA*) across COPD, CF, and healthy airway epithelium, with a consistent downregulation following flagellin treatment (**Supplementary File 1**), suggesting that TLR5 signaling operates similarly across these clinical conditions. This observation aligns with the study of Li *et al.* (25) that found that despite donor-specific transcriptional differences, flagellin-mediated signaling was similar in epithelium from healthy and chronic rhinosinusitis individuals.

TLR5 signaling is well characterized for initiating epithelial immune responses, driving the secretion of antimicrobial peptides and innate immune effector molecules, including chemokines, cytokines, and growth factors (4). This process orchestrates immune cell recruitment, activation, and modulation, shaping host defense against bacterial pathogens and maintaining mucosal immune homeostasis. The levels of *DEFB4A* mRNA (encoding beta-defensin 2) were significantly increased upon flagellin stimulation in healthy, COPD, and CF airway epithelia. While previous studies have demonstrated flagellin-induced *DEFB4A* upregulation in healthy bronchial epithelium (30), its expression has been reported to be downregulated in COPD explant lung samples (31). In our study, baseline *DEFB4A* levels were comparable across COPD, CF, and healthy airway epithelium (**Supplementary File 1**), suggesting that variations in tissue origin, COPD severity or time of analysis may account for these discrepancies. Flagellin also upregulated neutrophil-recruiting chemokines (*CXCL2/5/6/8*) in airway epithelium, irrespective of disease context, consistent with previous reports (18, 32, 33). Moreover, *CSF3*, encoding the granulocyte colony stimulating factor G-CSF emerged as a key marker of flagellin-induced immune activation, aligning with studies in mouse and porcine models (8, 13, 30, 34, 35). A recent study established a link between flagellin signaling, neutrophil recruitment, and therapeutic efficacy (13), likely mediated by chemokines and G-CSF, which promote neutrophil migration from the bloodstream and activation of antimicrobial neutrophils. Collectively, our findings demonstrate that COPD and CF airway epithelia retain a robust ability to enhance antibacterial defenses through antimicrobial peptide production and neutrophil-mediated immune activation.

The two cytokines IL-17C and IL-23, that are critical for innate immune responses (36, 37) were found to be prominently upregulated in airway epithelium by the treatment of flagellin at the air interface. This is consistent with previous observations that TLR signaling, particularly TLR5 signaling, impacts on the IL-17C/IL-23 axis in epithelium (37, 38). IL-17C is known to be rapidly upregulated in early stages of infection and is produced primarily by epithelial cells (36, 39–41). The specific receptor of IL-17C is the heterodimer IL-17RA/IL-17RE, and is expressed on both epithelial cells and some immune cells. Thus, IL-17C can play a key role in sustaining an autocrine loop in epithelial tissues, strengthening innate immune defenses through antimicrobial effects and recruitment of neutrophils. Interestingly, our study showed that *P. aeruginosa* infection followed by TLR5 stimulation is still effective at increasing *IL17C* expression in airway epithelium. Recently, epithelial-derived IL-23 has been implicated in regulation of immunity, as demonstrated in the *P. aeruginosa*-flagellin-TLR5-IL-23 axis linked to periodontitis in oral epithelial cells. (37). Moreover, Kim *et al.* showed comparable IL-23 expression in airway epithelial cells from healthy and diseased (COPD and COVID-19) lungs. Our study demonstrated that flagellin treatment increase by 10-fold *IL23A* expression across healthy, COPD, and CF airway epithelium, thereby reinforcing these findings. Together, the flagellin-driven upregulation of IL-17C and IL-23 highlights potential roles as immune modulators in airway epithelial defense mechanisms.

Infection of the airway epithelium creates a pro-inflammatory environment that can alter immune responsiveness (4, 26, 34, 42). Here, we demonstrate that despite the inflammation induced by the respiratory pathogens *P. aeruginosa* and *S. pneumoniae*, flagellin remains a potent stimulator of innate immune responses in human airway epithelium. Similarly, van Linge *et al.* reported that flagellin treatment enhanced the expression of antimicrobial genes (*S100A8, S100A9,* and *DEFB4A*) in human bronchial epithelial cells infected with a multidrug-resistant *Klebsiella pneumoniae* strain (24). Importantly, this flagellin-induced augmentation of immune responses in the context of infection has also been observed in murine and porcine models of bacterial respiratory infections that are not responding to antibiotic treatment (7, 8, 12, 35, 42). These findings collectively suggest that flagellin’s capacity to amplify antimicrobial defenses and cytokine production is not only preserved in an inflamed, infection-driven environment but also extends to clinically relevant pathogens associated with antimicrobial resistance and pneumonia. This reinforces the potential of flagellin as a promising immunomodulator for bacterial pneumonia, particularly in context where conventional antibiotic therapies are ineffective. This further supports the development of flagellin as an immunomodulator to treat bacterial pneumonia.

## Methods

### Flagellin

Recombinant flagellin FliC_∆174−400_ in Dulbecco’s Phosphate-Buffered Saline (PBS) or FLAMOD (FliC_Δ174-400_ harboring one extra N terminus amino acid in NaPi buffer: 10 mM phosphate, 145 mM NaCl, polysorbate 80 0.02 % (w/v) pH 6.5) were used (6–8, 12, 14).

### Epithelium culture and flagellin activation

Human airway epithelial cells were obtained from patients undergoing ectomy under informed consent and according to the declaration of Helsinki (Hong Kong amendment, 1989), with appropriate ethic approvals. MucilAir™ airway epithelium (Epithelix) from three donor sources (healthy individuals, COPD, and CF patients with *CFTR*<u>Δ*508* mutation</u>), were grown as described previously (43). Flagellin (0.1 or 0.7 µg, i.e., 0.3 or 2.1 µg/cm^2^) in 10 μl buffer or buffer alone was deposited apically after excess mucus removal.

### RNA extraction, sequencing and analysis

MucilAir™ inserts were lysed 4 h post-treatment with 150 μl RA1 buffer (Nucleospin^®^ RNA kit, Macherey-Nagel). RNA was extracted as per kit instructions. Libraries were prepared using the TruSeq Stranded mRNA kit (Illumina) and sequenced on the NextSeq500 system (Illumina). Differential expression analyses between flagellin- and PBS-treated conditions were performed using DESeq2. Gene set enrichment analysis (GSEA) was conducted on the ranked list of differentially expressed genes or DEG (R, ClusterProfiler version 4.6.2) with functions enrichGo and CompareCluster to identify Biological Processes. Analysis was carried out using Bioconductor (version 3.16.0). Benjamini-Hochberg method and threshold of 0.05 were used for False discovery Rate. The data have been deposited in NCBI’s Gene Expression Omnibus and are accessible through GEO Series accession number GSE304065.

### Bacterial strains and infections

*Streptococcus pneumoniae* serotype 19F (LILPNEUHC 19F) stock was prepared as described previously (8). For infection, 1 ml was thawed, washed with PBS, 0.1 ml diluted in 1.9 mL THYB, grown at 37°C for 3 h, diluted to 3×10^4^ CFU per 20 µl and added at the air-liquid interface. Treatment was followed apically 1 h later by flagellin/PBS, and amoxicillin (1.5 μg/ml) in basolateral medium. CFUs were determined in apical washes 24 h post-infection. Basolateral medium was collected for pro-inflammatory mediators. *Pseudomonas aeruginosa* (strain ATCC 9027) was grown on selective cetrimide agar plates at 37°C with 5% CO₂ for 24 h. For the inoculum, colonies were suspended in 2.2 mL of 0.9% NaCl, 1.25 mM CaCl₂, and 10 mM HEPES and adjusted to 50 CFU/20 µL. Epithelial cells were infected with 20 µL at the air-liquid interface, incubated at 37°C with 5% CO₂ and treated apically 16-17 h later with flagellin/PBS. Apical CFUs were determined 2 h post-treatment. Inserts were lysed for RNA extraction.

### RT-qPCR

RNA was reverse-transcribed with High-Capacity cDNA Archive Kit (Applied Biosystems) and cDNA amplified using SYBR-Green-based real-time PCR on a Quantstudio^TM^ 12K Real-Time PCR System (Thermo Fisher Scientific). Primers are listed in **Supplementary Table 1**.

### ELISA

Human β-defensin 2 and IL-8 were measured by commercial ELISA kits according to manufacturer’s instructions (Cusabio and Becton Dickinson Bioscience).

### Statistical Analysis

Results represent individual values and median or mean ± standard error of the mean (SEM). Mann-Whitney test (for two groups), or one-way analysis of variance (ANOVA) using GraphPad Prism software (version 8.2) with statistical significance set to p<0.05 were applied.

For extended materials and methods refer to supplementary materials

## Supporting information

List of DEG

List of GO

## Acknowledgements

We thank Mathilde Boissel and Julien Derop for the RNA-seq experiment design and analyses.

## Author contributions

Contribution: XL, DC, MCC, CCC, MB, TvdP, and SC planned studies, performed experiments and analyzed data. YZ, ISG, AB and MD performed and analyzed the RNA-seq experiments. XL, CC, MB, TvdP, SC, and JCS wrote the paper with input from all authors. SC and JCS supervised the project.

