## Supplementary Materials for "Flagellin-mediated TLR5 activation enhances innate immune responses in healthy and diseased human airway epithelium"

The supplementary materials include:

- Supplementary methods
- 1 supplementary table
- 3 supplementary figures
- 2 supplementary files (as separate files that list the differentially expressed genes, and the associated biological processes using gene ontology, respectively)

### Supplementary methods

#### Flagellin production

The recombinant flagellin $FliC_{\Delta174-400}$ (derived from *Salmonella enterica* serovar Typhimurium FliC) was produced with a histidine tag, as described previously (1-3). The protein $FliC_{\Delta174-400}$ (batch 04052017, 2 mg/mL; excipient Dulbecco's Phosphate-Buffered Saline (PBS, Gibco, Grand Island, NY)) was certified to be biologically active in reporter cells (HEK-Dual™ hTLR5 cells assay; Invivogen) and in mouse assays, and LPS contamination by limulus assay (Associates of Cape Cod Inc., East Falmouth, MA) was estimated < 20 pg of LPS per µg of flagellin. In some experiments, FLAMOD (recombinant flagellin $FliC_{\Delta174-400}$ harboring one extra amino acid at the N terminus) resuspended in NaPi buffer: 10 mM phosphate, 145 mM NaCl, polysorbate 80 0.02 % (w/v) pH 6.5 was used as described previously (4, 5).

#### Epithelium culture and flagellin activation

The MucilAir™ (Epithelix, Plan-les- Ouates, Switzerland) model are derived from primary human airway epithelial cells and are composed of basal cells (progenitor), goblet cells (mucus producing), and ciliated cells. The MucilAir™ model were differentiated *in vitro* on inserts at the air-liquid interface. Cells were grown in a humidified incubator at 37 °C with 5% $CO_2$ in the MucilAir™ Culture Medium supplemented with 1% penicillin-streptomycin (Gibco BRL, Rockville, USA) with basolateral medium changed every other day as described previously (6). On the day of the experiment, cells were washed with 37°C PBS to remove any excess of mucus. Three sources of donors were used: healthy individuals, patients with COPD, and patients with cystic fibrosis (CF) harboring the mutation $CFTR^{\Delta F508}$. To mimic airway administration, flagellin (0.1 or 0.7 µg, i.e., 0.3 or 2.1 µg/cm$^2$) in 10 µl buffer was deposited on the apical compartment of epithelium. Vehicle buffer (PBS or NaPi) deposition was used as control.

#### Bacterial strains and infections

*Streptococcus pneumoniae* serotype 19F (clinical isolate LILPNEUHC 19F) was obtained from the University Hospital of Lille (Frédéric Wallet, CHU Lille, France) and working stocks were prepared as described previously (5). Briefly, fresh colonies grown on blood-agar plates were incubated in Todd Hewitt Yeast Broth (THYB, Sigma-Aldrich, Saint-Louis, MO) at 37°C with 5% $CO_2$ until the OD at 600 nm reached 0.7–0.9 units. Cultures were stored at −80°C in THYB + glycerol 12% (vol./vol.) for up to 3 months. For infection, 1 ml working stocks were thawed and washed with PBS, 0.1 ml were then diluted in 1.9 mL THYB and grown at 37°C for 3 h. and diluted to the appropriate concentration. Bacteria (as $3\times10^4$ CFU in 20 µl) were added at the air interface, treated 1 h after bacterial inoculation by adding at the air interface flagellin or PBS in 10 µl and in the basal medium, amoxicillin at 1.5 µg/ml. Twenty-four hours post-infection, bacteria number were determined in apical washes and the basolateral medium was collected to evaluate the production of pro-inflammatory mediators using ELISA assays. The *Pseudomonas aeruginosa* ATCC 9027 strain was initially isolated from a patient with otitis and was grown on the selective cetrimide agar plates (Biomerieux SA, Marcy-L'Etoile, France). A bacterial stock was prepared from a single colony transferred into a cryovial containing cryopreservation beads and medium (Dutscher SAS, France) and then stored at -80°C. To prepare the inoculum, cetrimide agar were inoculated with a bead and incubated at 37°C with 5% $CO_2$ for 24 h. Bacterial colonies were suspended in 2.2 mL of a solution containing 0.9% NaCl, 1.25 mM $CaCl_2$, and 10 mM HEPES. The $OD_{600}$ was measured to calibrate the inoculum and then validated by CFU counts on agar plates. The infection of MucilAir™ cells was carried out by applying at the air-liquid interface 20 µL containing 50 CFU. The infection was then continued by incubating the cells at 37°C with 5% $CO_2$. Cells were then treated with flagellin or PBS in 10 µl at the air interface 16 h or 17 h after bacterial inoculation. Two hours post-treatment, the number of CFUs in the apical compartment was determined and the cells were lysed for RNA extraction and gene expression analysis.

**Analysis of gene expression by real time quantitative PCR (RT-qPCR)**

Total RNA was reverse-transcribed with the High-Capacity cDNA Archive Kit (Applied Biosystems, Foster City, CA, USA). The cDNA was amplified using SYBR-Green-based real-time PCR on a Quantstudio™ 12K Real-Time PCR System (Thermo Fisher Scientific, Carlsbad, CA, USA). Specific primers used are listed in **Supplementary Table 1**. Relative mRNA levels were determined by comparing the PCR cycle threshold (Ct) for the gene of interest to the reference genes *ACTB, B2M, and H3.3* (ΔCt) and then the ΔCt values for flagellin-treated vs PBS-treated groups (ΔΔCt). Ct upper limit was fixed to 35 cycles.

**Supplementary Table 1. List of primers used for transcriptional studies**

| Target genes | Forward primer | Reverse primer |
|---|---|---|
| *ACTB* | ATTGGCAATGAGCGGTTC | CGTGGATGCCACAGGACT |
| *B2M* | TTCTGGCCTGGAGGCTATC | TCAGGAAATTTGACTTTCCATTC |
| *CCL4* | CTTCCTCGCAACTTTGTGGT | CAGCACAGACTTGCTTGCTT |
| *CCL20* | CCAAGAGTTTGCTCCTGGCT | TGCTTGCTGCTTCTGATTCG |
| *CCL22* | GTGGCGCTTCAAGCAACT | AGACGGTAACGGACGTAATCA |
| *CLDN24* | GGAGGGTGTCTGCTCAACTG | CATAGTGGCCCAAAGCTAGG |
| *CSF3* | GTGCTGCTCGGACACTCTCT | GAAAAGGCCGCTATGGAGTT |
| *CXCL2* | CCCATGGTTAAGAAAATCATCG | CTTCAGGAACAGCCACCAAT |
| *CXCL3* | AAATCATCGAAAAGATACTGAACAAG | GGTAAGGGCAGGGACCAC |
| *CXCL6* | GTCCTTCGGGCTCCTTGT | CAGCACAGCAGAGACAGGAC |
| *CXCL8* | CACCGGAAGGAACCATCTCA | GGAAGGCTGCCAGAGAGC |
| *DEFB4A* | TCAGCCATGAGGGTCTTGTA | AGGATCGCCTATACCACCAA |
| *H3* | AGACTGCCCGCAAATCGAC | CTTGCGAGCGGCTTTTGTA |
| *IDO1* | CAAAGCAGCGTCTTTCAGTG | AGGAACTGAGCAGCATGTCC |
| *IL1B* | TACCTGTCCTGCGTGTTGAA | TCTTTGGGTAATTTTTGGGATCT |
| *IL17C* | CCCTCAGCTACGACCCAGT | CTTCTGTGGATAGCGGTCCT |
| *IL23A* | TGTTCCCCATATCCAGTGTG | TCCTTTGCAAGCAGAACTGA |
| *MUC2* | ACAAGGACTGCACCCCATC | AACACGCAGGCATCGTAGTA |
| *MUC5AC* | GGGACAAGAAGACCAGCATC | CGTCGAAGTTCCCACACAG |
| *RELN* | TGCTGGAATACACTAAGGATGC | GAAGGCACTGGGTCTGTACG |
| *SMIM11B* | GGACACCTGTCATCGCTTCT | GCCCCCAACAGACAAGTACA |
| *SPRR2* | TGGTACTTGAGCACTGATCTGC | TGCACTGCTGCTGTTGATAA |
| *TACR1* | GGTCTACCTGGCCATCATGT | GAAGCCCAGACGGAACCT |
| *TFF1* | CCCCTGGTGCTTCTATCCTAA | GATCCCTGCAGAAGTGTCTAAAA |
| *TNFAIP6* | GGCCATCTCGCAACTTACA | GCAGCACAGACATGAAATCC |

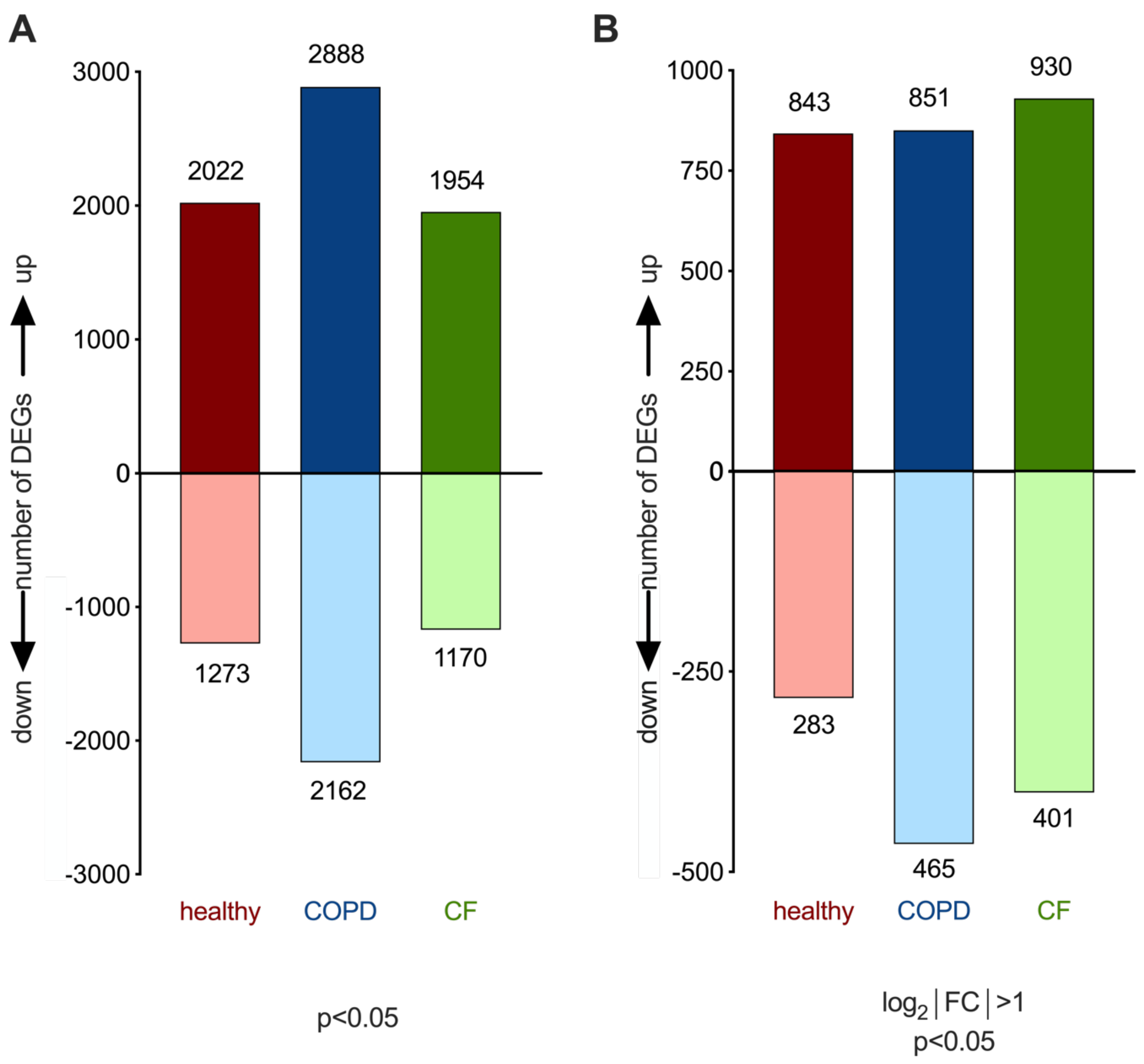


**Supplementary Figure 1. TLR5 signaling induces transcriptional activation of genes in airway epithelium derived from healthy individuals and COPD and CF patients.** Flagellin FliC$_{\Delta 174\text{-}400}$ (0.7 µg in 10 µL) or the diluent solution (PBS) was applied for 4 h at the air interface of MucilAir™ cells from healthy donors, COPD or CF patients. RNA was processed for RNA-seq analysis. Analysis of the pattern of transcriptional activation was conducted to define the number of genes that were up-regulated and down-regulated by flagellin treatment. (**A**) Number of differentially expressed genes (DEG) in various clinical situations based on the *p* value ($p<0.05$). (**B**) Number of DEG based on the *p* value ($p<0.05$) and a fold change (FC) in expression >2 or <0.5.

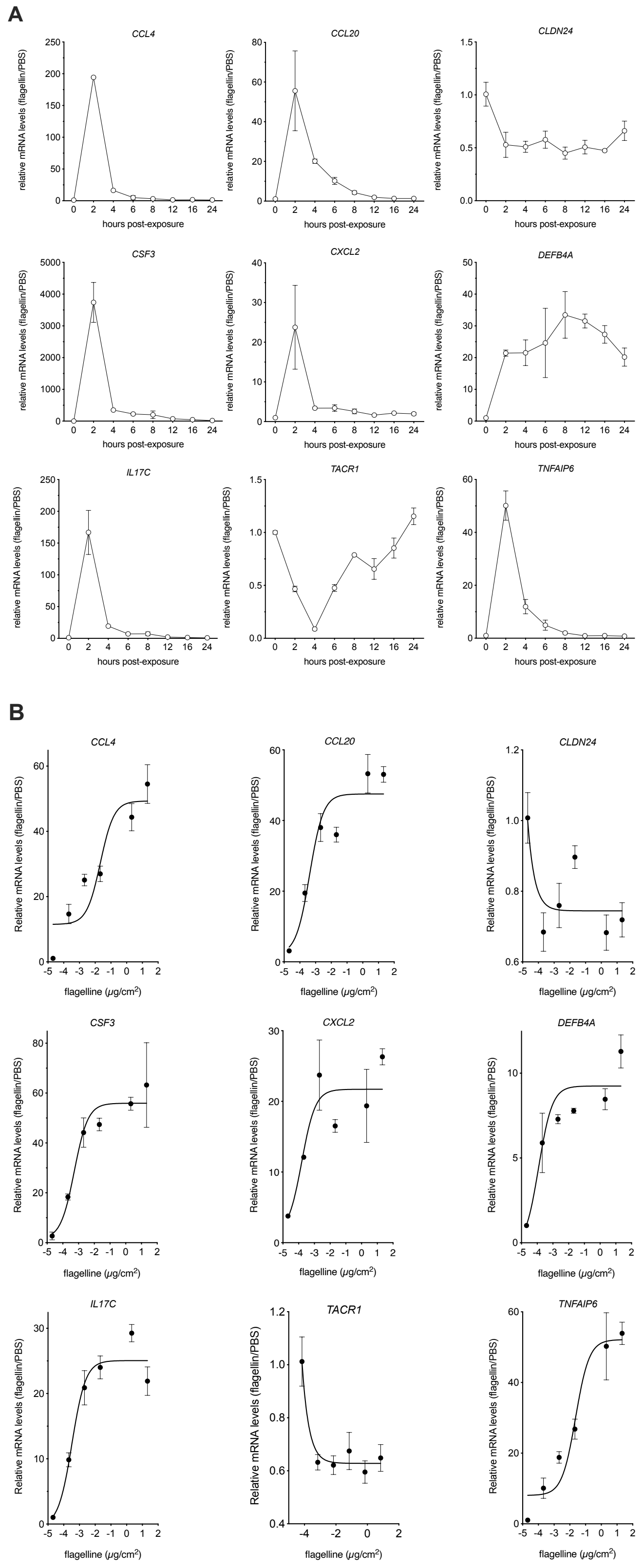


**Supplementary Figure 2. TLR5 signaling in airway epithelium is regulated temporally and in a flagellin-dose dependent manner.** Flagellin FliC$_{\Delta174\text{-}400}$ (indicated amounts in 10 µL) or the diluent solution (PBS) was applied at the air interface of MucilAir™ cells from healthy donors. RNA was processed for RT-qPCR analysis at indicated times. (**A**) Kinetic analysis of gene expression after stimulation with 0.7 µg flagellin, i.e., 2.1 µg/cm$^2$ (n=2 per time). (**B**) Gene expression analysis 2 h post-exposure to various amounts of flagellin from 0.000021 to 21 µg/cm$^2$ (n=4 per dose). mRNA levels were normalized to house-keeping genes and the condition treated with PBS was set arbitrarily at the value of 1. A nonlinear fitting of data was performed using a log(agonist) vs response curve with three parameters. The ED50 can be estimated as the concentration of agonist that produces 50% of the maximal biological response, i.e., gene expression level.

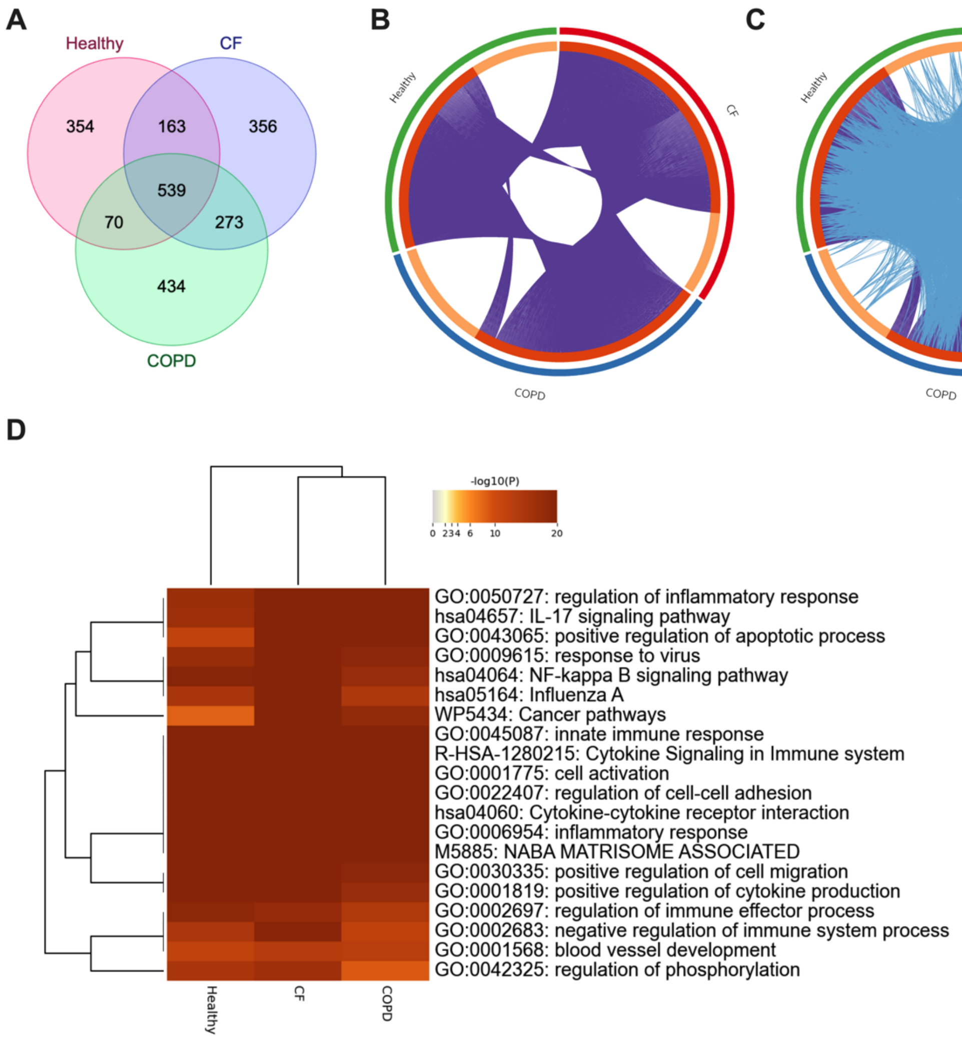


**Supplementary Figure 3. Flagellin results in high overlap of common genes and functions between healthy, COPD and CF conditions.** Flagellin FliC$_{\Delta 174\text{-}400}$ (0.7 µg in 10 µL) or the diluent solution (PBS) was applied at the air interface of MucilAir™ cells from healthy individuals, COPD patients or CF patients harboring the *CFTR*<u>*Δ508*</u> mutation. After 4 h incubation, RNA was processed for RNA-seq analysis (n=3-4 per condition). Total RNA was extracted and processed for RNA sequencing. Metascape (https://metascape.org; (7)) was used for gene overlap analysis. (**A**) Venn diagram illustrating the number of unique and overlapping genes across different conditions. (**B**) Circos plot depicting gene overlap among healthy, COPD, and CF conditions. Outer arcs represent each condition, dark orange inner arcs indicate shared genes connected by purple lines, and light orange arcs denote unique genes for each condition. (**C**) The same circus plot from (**B**) with an additional layer representing the extent of functional overlap among gene lists based on shared ontology terms (indicated by blue lines). (**D**) Dendogram of enrichment ontology clusters across the three conditions. The heatmap cells are colored by their *p*-values.
